# Quantitative and molecular differences distinguish adult human medullary and extramedullary haematopoietic stem and progenitor cell landscapes

**DOI:** 10.1101/2020.01.26.919753

**Authors:** Nicole Mende, Hugo P Bastos, Antonella Santoro, Kendig Sham, Krishnaa T. Mahbubani, Abbie Curd, Hitoshi Takizawa, Nicola K Wilson, Bertie Göttgens, Kourosh Saeb-Parsy, Elisa Laurenti

**Author notes:** equal contribution (Nicole Mende, Hugo P Bastos).

## Abstract

In adults, the bone marrow (BM) is the main site of haematopoietic stem and progenitor cells (HSPCs) maintenance and differentiation. It is known that other anatomical sites can contribute significantly to blood production under stress conditions. However limited tissue availability restricts our knowledge on the cellular, molecular and functional composition of extramedullary HSPC pools in humans at steady state or under stress. Here we describe the landscape of human HSPC differentiation across the three major haematopoietic anatomical sites: BM, spleen and peripheral blood (PB), using matched tissues isolated from the same individuals. Single cell RNA-seq of 30,000 HSPCs and 700 phenotypic haematopoietic stem cells and multipotent progenitors (HSC/MPP) demonstrates significantly different dynamics of haematopoiesis between BM and extramedullary tissues. Lineage-committed progenitors of spleen and PB do not actively divide, whereas BM is the primary site of progenitor proliferation. The balance of differentiation in spleen and PB is skewed towards the lymphoid and erythroid lineages, whereas in BM it is tilted towards megakaryocytic and myeloid progenitors. Extramedullary tissues also harbour a molecularly defined subset of HSC/MPP not found in the BM, which is marked by a specific acto-myosin cytoskeletal signature and transcriptional priming for division and lineage differentiation. Collectively, our findings define a unique cellular and molecular structure of the haematopoietic landscape in extramedullary organs, positioned for rapid lineage-primed demand-adapted haematopoiesis. These data also provide a framework for better understanding of human extramedullary haematopoiesis in health and disease.

## INTRODUCTION

In adults, more than 99% of haematopoietic stem and progenitor cells (HSPCs) reside within the bone marrow (BM) (Nombela-Arrieta and Manz, 2017; Soderdahl et al., 1998), giving rise to all mature blood cells. However, a small proportion of HSPCs constitutively migrate through peripheral blood (PB), probably to repopulate injured areas of BM or to assist local production of immune cells in extramedullary tissues, such as spleen, lung and liver (Lefrançais et al., 2017; Massberg et al., 2007; Wright et al., 2001). In adulthood, HSPC migration and differentiation outside the BM, also called extramedullary haematopoiesis (EMH), is mostly associated with haematopoietic stress. The spleen, for example, was the first organ reported to contain stem and progenitor cells that are able to rescue irradiation-induced haematopoietic failure (Jacobson et al., 1951). Today, the splenic red pulp is recognized as one of the most common sites for EMH in response to haematopoietic challenges like anaemia (Harandi et al., 2010; Lenox et al., 2005; Perry et al., 2007), increased demand of blood cells during pregnancy (Inra et al., 2015; *Maymon et al., 2006*; Nakada et al., 2014; Oguro et al., 2017); and various pathologies, such as myeloproliferative disorders (O’Malley et al., 2005) and chronic inflammation/infection (Griseri et al., 2012; Masuya et al., 2014). Mechanistically, upon BM injury or stress, BM HSPCs traffic to the spleen and are attracted to specific stromal cells producing SCF and CXCL12 (Inra et al., 2015; Miwa et al., 2013; Wang et al., 2015).

At steady-state in mice, the spleen contains a rare population of phenotypic HSCs, which, despite being 15-times less frequent than in BM, possesses similar long-term repopulating capacity as their BM counterparts (Morita et al., 2011). Similarly, human spleen and BM phenotypic HSPCs did differ in frequency, but not in long-term culture initiating cell capacity (Dor et al., 2006). However, whether human spleen contains any cells with long-term *in vivo* repopulation capacity has not been tested. Interestingly, in mice, most phenotypic HSCs from BM and spleen were not replaced by circulating HSCs after 14 weeks of parabiosis (Morita et al., 2011). This suggests that extramedullary HSPCs are long-term residents of their respective tissues, and not merely BM-derived circulating HSPCs transiently occupying a distinct niche. In addition, progenitor cell types exclusively residing in the spleen have been identified in mice, indicating that splenic HSPCs might also play a distinctive, yet unclear role, in steady-state haematopoiesis. These studies in mice revealed a myelo-erythroid-restricted population different from any BM progenitor (Mumau et al., 2017), and an erythroid-restricted cell type, termed stress BFU-E, which has been shown to expand and gain some level of self-renewal in response to anaemia or injury (Harandi et al., 2010; Perry et al., 2007). Altogether these studies point to a role of resident splenic HSPCs in adult blood cell production. However, because of limited tissue availability from healthy donors, extremely little is known about the human spleen HSPC landscape at steady state and its contribution to undisturbed haematopoiesis.

Here we employed single cell RNA-seq (scRNA-seq) and functional assays to compare the molecular and functional composition of the human HSPC pool in BM, PB and spleen, and gain insights into human extramedullary haematopoiesis. We find that the structure of the human HSPC hierarchy in spleen and PB clearly differs from that of BM through the proportions of lineage-committed and multipotent progenitors (MPP). In addition, we identify a molecularly defined HSC/MPP subtype present exclusively in extramedullary tissues, which is marked by the expression of specific cytoskeleton genes and possesses all the hallmarks of a quiescent cell poised to rapidly respond to stress.

## RESULTS

### Construction of a multi-tissue single-cell transcriptomic map of adult human HSPC differentiation

Comprehensive characterization of the adult human HSPC transcriptional landscape across hematopoietic tissues requires analysis of matched tissues harvested from the same individuals. Access to adult human tissues is scarce and samples from deceased organ donors are thus a valuable source for the study of adult human biology. To date, organ donors have been used to chart the cellular composition and physiology of adult human tissues (Braga et al., 2019; MacParland et al., 2018; Reyfman et al., 2018), under what in most cases remains the best approximation of a steady-state healthy situation. Here we used HSPCs from matched BM, PB and spleen of 2 deceased organ donors with no clinical signs of acute infection or known history of chronic disease (age: 19 and 35 years old, other characteristics in **Table S1)**. A droplet approach yielded transcriptomic data of a total of 30,873 CD19^-^ CD34^+^ HSPCs across all three tissues (13,393 from donor 1 and 17,481 from donor 2). In parallel, the transcriptomes of 697 single HSC/MPPs from the same donors and tissues were determined using the Smart-seq2 protocol (**Figure 1A,** see methods). These were isolated as single CD19- CD34+ CD38-CD45RA-cells, a fraction highly enriched for multipotent cells with long and short-term reconstitution potential (Majeti et al., 2007; Notta et al., 2011) (hereby referred to as HSC/MPPs, **Figure S1A**).

**Figure 1.**
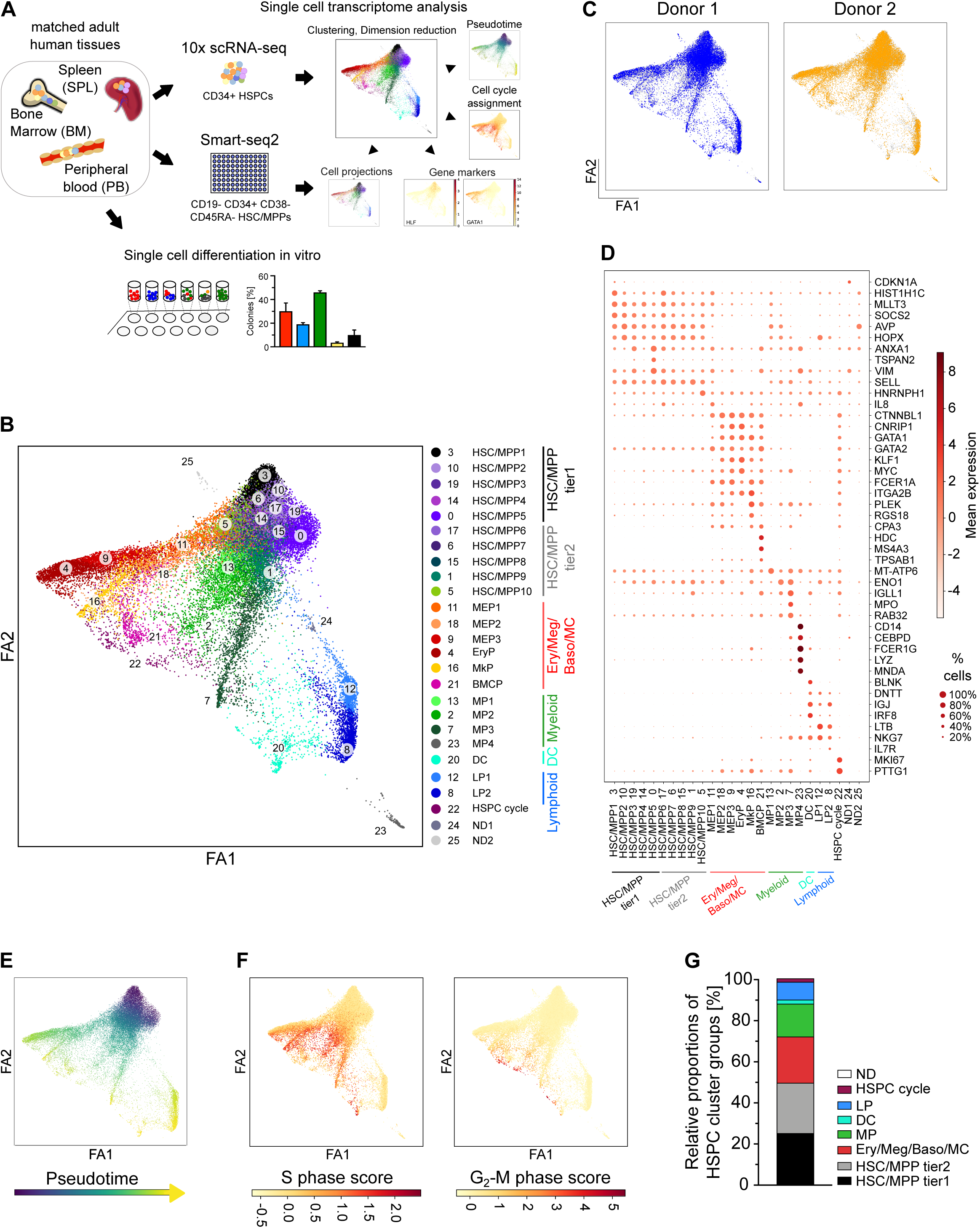
Single cell transcriptomic landscape of HSPC across medullary and human adult haematopoietic tissues. **A)** Outline of experimental approach. CD19- CD34+ HSPCs were isolated from matched bone marrow (BM), peripheral blood (PB) and spleen (SPL) from two different deceased adult organ donors. scRNA-seq was performed using the commercial 10x Genomics approach. In parallel phenotypic HSC/MPPs were purified by flow cytometry and processed using the Smart-seq2 protocol. Single-cell RNAseq data was used to compute a force-directed graph (FDG) representing the transcriptomic landscape of adult human HSPCs across all three organs. Functional features of phenotypic HSC/MPPs from the same tissues but different donors were also assessed *in vitro*. **B-G)** Analysis of 10x Genomics scRNAseq data from 30,873 cells, combining all tissues and both donors. **B)** Force-directed graph of the multisite HSPC landscape. Colours and numbers indicate clusters of transcriptionally distinct cells as defined by Louvain clustering. Clusters were annotated using known lineage and stem cell marker genes found amongst the most differentially expressed genes in each cluster (**Table S4**). In addition, clusters were grouped into 6 main HSPC groups as indicated on the right: HSC/MPP tier1, HSC/MPP tier2, Ery/Meg/Baso/MC, Myeloid progenitors, DC progenitors and lymphoid progenitors. ND: clusters which identity could not be defined using known marker genes. HSC/MPP: haematopoietic stem cell/multipotent progenitor; MEP: Megakaryocyte-erythroid progenitor; EryP: erythroid progenitor; MkP: megakaryocytic progenitor; BMCP: Basophil/mast cell progenitor; MP: myeloid progenitor; DC: dendritic cell progenitor; LP: lymphoid progenitor. **C)** FDG of the same data as in (B) showing the distribution of cells from individual donors across the FDG space in separate colours. **D)** Expression values of selected marker genes for all Louvain clusters (top 100 in **Table S4**). Circle colour shows mean scaled expression values and circle size represents the proportion of expressing cells per cluster. **E)** Pseudotime was assigned to each cell of the HSPC landscape choosing a cell in cluster 3 as a starting point. Each cell of the FDG is coloured by the rank of its pseudotime score. **F)** FDGs show single cells coloured by their assigned S-phase score (left) and G_2_-M phase score (right) as calculated by the cell phase scoring method defined in (Satija et al., 2015). **G)** Cluster composition using the groups as defined in (B). HSPCs from both donors and all tissues are combined in this graph, data from individual donors is shown in **Figure S1D**.

To generate a reference map of all analysed haematopoietic tissues and both donors (**Table S2** for the number of cells passing quality control) we combined all cells from both datasets using the Seurat alignment method (Stuart et al., 2019). The Louvain clustering algorithm (Blondel et al., 2008) applied to single cells from all organs and both donors partitioned HSPCs into 26 transcriptionally distinct cell clusters **(Figure 1B, Figure S1B, Table S3).** A force-directed graph (FDG) layout revealed a fork-like cluster organisation **(Figure1B,** UMAP in **Figure S1B),** with a core of 10 clusters grouped together at the head of the fork and all residual clusters spreading across three main branches. Cells from both donors were present in all clusters (**Table S3**) and similarly distributed across the HSPC landscape **(Figure 1C, Figure S1C)**. Analysis of highly expressed marker genes for each cluster identified that the clusters at the head of the fork (clusters 3,10,19,14,0,17,6,15,1 and 5) are marked by stem cell-associated genes such as *SOCS2, MLLT3* and *HOPX* (Calvanese et al., 2019; Nguyen et al., 2019; Vitali et al., 2015; Zhou et al., 2015) **(Figure 1D, Table S4**). Clusters within the three branches could be attributed to the main haematopoietic lineages. Consistent with other single cell studies of human HSPCs (Hay et al., 2018; Pellin et al., 2019; Popescu et al., 2019), we observed grouping of erythroid (Ery) progenitors (marked by high expression of *GATA1, FCER1A* and *CTNNBL1),* megakaryocytic (Meg) progenitors (high expression of *PLEK* and *ITGA2B*) and basophil/mast cell (Baso/MC) progenitors (high transcript levels of *HDC, MS4A3* and *TPSAB)* in one branch of the landscape. The second branch contained myeloid (My) progenitors marked by *ENO1*, *MPO* and *CEBPD* and the last branch corresponded to lymphoid (Ly) clusters (identified by high expression levels of *IGJ, LTB* and *NKG7*). We also observed a cluster of dendritic cell progenitors (DCs, cluster 20) with transcriptional characteristics of plasmacytoid dendritic cells, such as high expression of IRF7, IRF8, IGJ and BLNK.

Projection of Smart-seq2 transcriptomic data from phenotypic HSC/MPPs that were purified by flow cytometry from the same donors onto their respective HSPC landscape confirmed that the top of the fork contained most immature stem and progenitor cells (76.6% all projected phenotypic HSC/MPP) **(Table S5)**. Pseudotime analysis (Haghverdi et al., 2016) confirmed this observation as the most immature cells of the haematopoietic hierarchy were earlier along the pseudotime axis. **(Figure 1E)**. Cell cycle assignment showed that, with the exception of the Ly clusters (clusters 8 and 12), the top of the fork clusters had the lowest proportion of cells in the S-G_2_-M cell cycle phases **(Figure 1F, Table S6)**. 5 clusters (3, 10, 19, 14 and 0) had 15% or less cells in S-G_2_-M and also contained the highest proportions of projected phenotypic HSC/MPPs. These were grouped into an ‘HSC/MPP tier1’ category, and were considered to be the most immature cells in the dataset. Clusters with 15-25% S-G_2_-M cells were classified as ‘HSC/MPP tier2’ (clusters 17, 6, 15, 1 and 5). Cell cycle activity markedly increased along the My and Ery/Meg/Baso/MC branches of the force-directed graph, with the most actively cycling progenitors at their tips **(Table S6)**. On the basis of known literature, top marker genes of each cluster and cell cycle analysis we assigned cluster nomenclature as listed in **Figure 1B**. With this nomenclature, the adult human haematopoietic landscape of HSPCs combining the three main haematopoietic tissues (BM, PB and spleen) of both donors comprises about 25% Tier1 HSC/MPPs, 25% Tier2 HSC/MPPs, 16 % My-committed progenitors, 11 % Ly- and dendritic cells (DC) committed progenitors and 22 % Ery/Meg/Baso/MC progenitors **(Figure 1G,** composition for each donor in **Figure S1D)**. Similar cluster identities and proportions were found if the cells from both datasets were combined with Scanorama instead of the Seurat alignment method (Hie et al., 2019) (**Figure S1E-G**). The general structure of the HSPC landscape outlined here overall resembles that described for human HSPC hierarchies in foetal, neonatal and adult life (Hay et al., 2018; Pellin et al., 2019; Popescu et al., 2019; Velten et al., 2017; Zheng et al., 2018), and provides a reference framework on which to compare the distinct organs.

### BM is the primary site of proliferation for lineage-committed progenitors

To investigate if distinct cell types identified above were equally represented in each haematopoietic tissue, we calculated the frequency of each cluster in each organ combining both donors. Whereas all clusters were present in all tissues, their frequencies varied between tissues (**Figure 2A, Table S3** and individual donors in **Figure S2A**). To highlight clusters specifically enriched in extramedullary organs, we computed a ‘location score’ for PB and spleen (equivalent to the fraction of cells in each extramedullary tissue compared to the sum of cells in BM and that tissue; see methods) **(Figure 2B)**. This indicated enrichment of specific Tier1 and Tier2 HSC/MPP clusters in spleen and PB. Within the lineage committed progenitor branches, both extramedullary tissues contained significantly less My and DC progenitors, but an increased proportion of non-dividing Ly-committed progenitor cell types committed to B, T and NK lineages **(Figure 2B-C, Figure S2B**). The overall size of the Ery/Meg/Baso/MC branch was only slightly increased in PB and not significantly different in spleen compared to BM (**Figure 2B-C, Figure S2B**).

**Figure 2.**
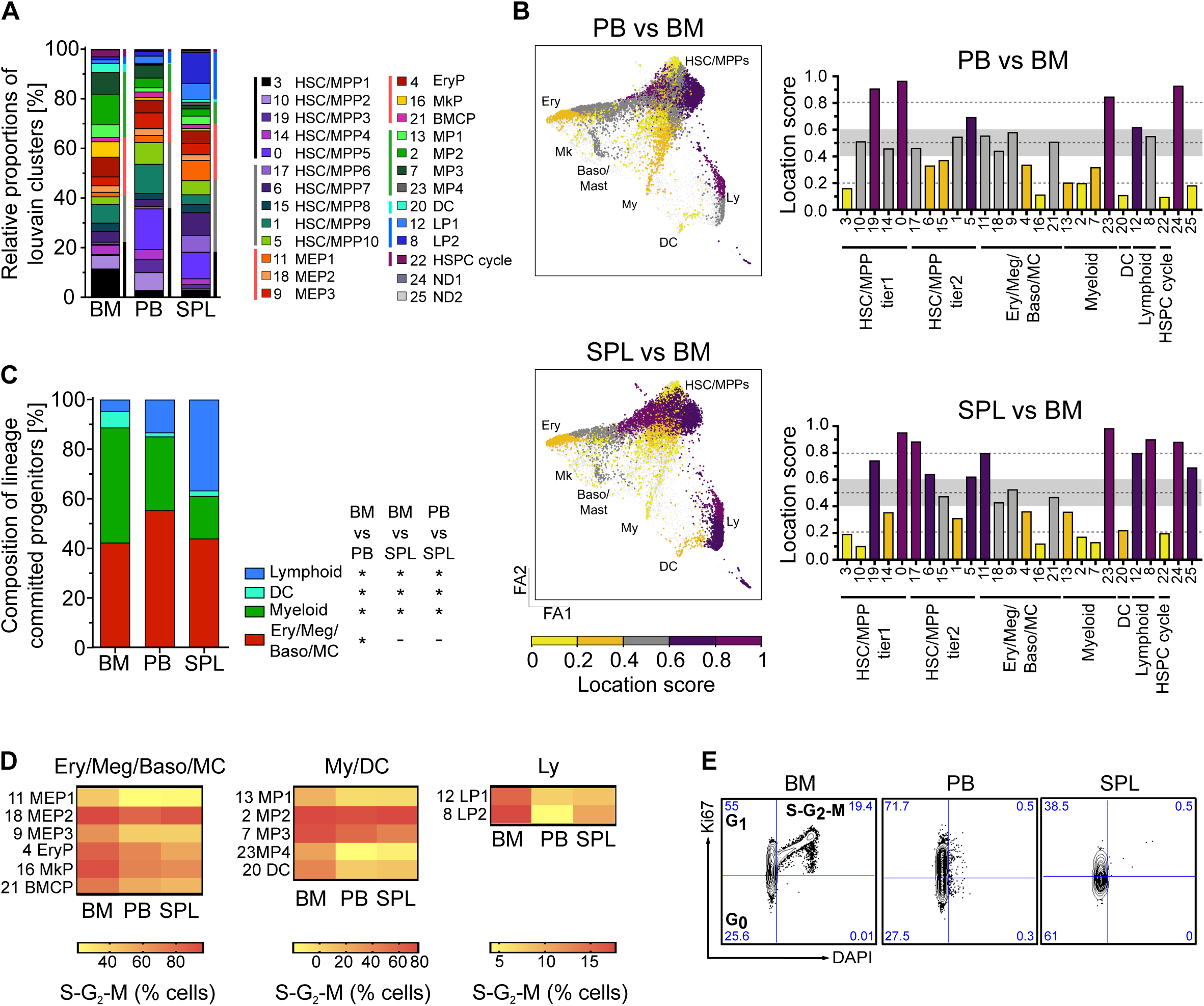
Comparison of relative cell type abundances and cell cycle properties in BM, PB and spleen. **A-D)** Analysis of 10x Genomics scRNAseq data from 30,873 cells, combining both donors but comparing different tissues. **A)** Relative proportion of each cluster within the HSPC landscape of the respective haematopoietic tissue. Donor 1 and 2 were combined for this analysis. Individual donors are shown in **Figure S2A**. **B)** A location score was computed to compare the size of each cluster in PB and spleen to the cluster sizes in BM as reference organ. The location score, for any given cluster, was calculated as: (number cells in that cluster for spleen or PB)/ (number of cells in BM for that cluster + number of cells in spleen or PB for that cluster). FDGs show each cluster along the HSPC landscape coloured by its location score. The score for each cluster is also shown in the bar graphs to the right of each FDG. A score >0.6 (purple) indicates higher representation of the cluster in PB or spleen compared to BM, a score <0.4 (yellow) indicates that this cluster is enriched in BM compared to spleen or PB. Clusters that were equally represented across both organs have a score between 0.4 and 0.6 (grey). **C)** Bar graph showing the relative composition of lineage-committed progenitors in BM, PB and spleen. Each branch was defined as shown in Figure 1B (Lymphoid: clusters 8, 12; DC: cluster 20; Myeloid: clusters 2, 7, 13, 23; Ery/Meg/Baso/MC: clusters 4, 9, 11, 16, 18, 21). Both donors were combined for this analysis, data of individual donors is shown in **Figure S2B**. Fisher test. * p<10^-5^. **D)** Heatmaps show the average fraction of cells in S-G_2_-M phase for each lineage-committed progenitor cluster as assigned by the cell phase scoring algorithm as described by (Satija et al., 2015). **Table S6** reports raw values. **E)** Flow cytometry plots of BM (left), PB (middle) and spleen (right) CD19^-^ CD34^+^ CD38^+^ progenitor cells in G_0_ (Ki-67^-^DAPI^-^), G_1_ (Ki-67^+^DAPI^-^) and S-G_2_-M (Ki-67^+^DAPI^+^) cell cycle phases. % of cells in each quadrant is indicated. SPL: spleen.

We next performed transcriptome-based cell cycle assignment (Satija et al., 2015) of the clusters in the lineage-committed progenitor branches of different tissues. Strikingly, all BM clusters displayed increased proportions of cycling cells (S-G_2_-M phases) than in PB or spleen (in most cases >2-fold increase, **Figure 2D, Table S6).** This highly proliferative phenotype of BM progenitors was confirmed by Ki67/DAPI flow cytometry of CD19- CD34+ CD38+ cells from independent donors, where cells in S-G_2_-M were only observed in BM **(Figure 2E)**. In addition, a large proportion of spleen and PB progenitor cells were found in G_1_ suggesting that they may be in a poised state, ready for rapid response to extrinsic stimuli. Overall, our data suggest that BM is the primary site of active proliferation and differentiation during human steady-state haematopoiesis.

### The balance of Ery/Meg priming differs between BM and SPL

Given the documented role of spleen-resident Ery progenitors in stress erythropoiesis in mice (Harandi et al., 2010; Lenox et al., 2005; Perry et al., 2007), we reasoned that analysis of the Ery/Meg/Baso/MC branch may provide insights into potential contributions of HSPCs from distinct sites to steady-state and stress haematopoiesis. Differentiation along the Ery/Meg/Baso/MC branch involves consecutive oligopotent steps (here represented by the megakaryocyte erythroid progenitor (MEP) clusters MEP1 (cluster 11), MEP2 (cluster 18) and MEP3 (cluster 9), named based on their position along the FDG derived trajectory), followed by unilineage progenitors, which are actively dividing (**Figure 1F, 2D**). To better map Ery or Meg priming along this branch, we computed a score of genes shown to be up-regulated during human Ery or Meg differentiation (Lu et al., 2018; Psaila et al., 2016). This analysis revealed that across all tissues, whereas expression of Ery-associated genes gradually increased from MEP1 to erythroid progenitors (EryP, cluster 4), expression of Meg-associated genes was highest in MkP and MEP1 (**Figure 3A**). We concluded that MEP1 is primed to both Meg and Ery fates, but more Meg-primed than other MEP subsets. In contrast, MEP2 and 3 are predominantly precursors of Ery progenitors. Despite the Ery/Meg/Baso/MC branch being overall equally represented in BM and extramedullary tissues (**Figure 2C**), there were marked differences in the abundance of the individual clusters within the branch **(Figure 3B**, for data of individual donors see **Figure S3A)**. The relative size of Baso/MC progenitors (BMCPs) did not differ between organs. However, megakaryocyte progenitors (MkP, cluster 16) were significantly more abundant in BM HSPCs (approximately 6-fold compared to spleen and PB, **Figure 3B, S3A**). At the molecular level, we observed increased Meg-priming in the BM Ery/Meg/Baso/MC branch, as evidenced by significantly higher Meg scores not only in BM MkP, but also early on in BM MEP1 compared to spleen (**Figure 3C**). Consistently Meg-associated genes such as *ITGA2B, SELP, RGS18*, and *NFIB* were expressed at significantly higher levels in MEP1 of BM than spleen (**Figure 3D, Table S7**). These data suggest that the whole route of Meg differentiation is enhanced in BM HSPCs compared to spleen.

**Figure 3.**
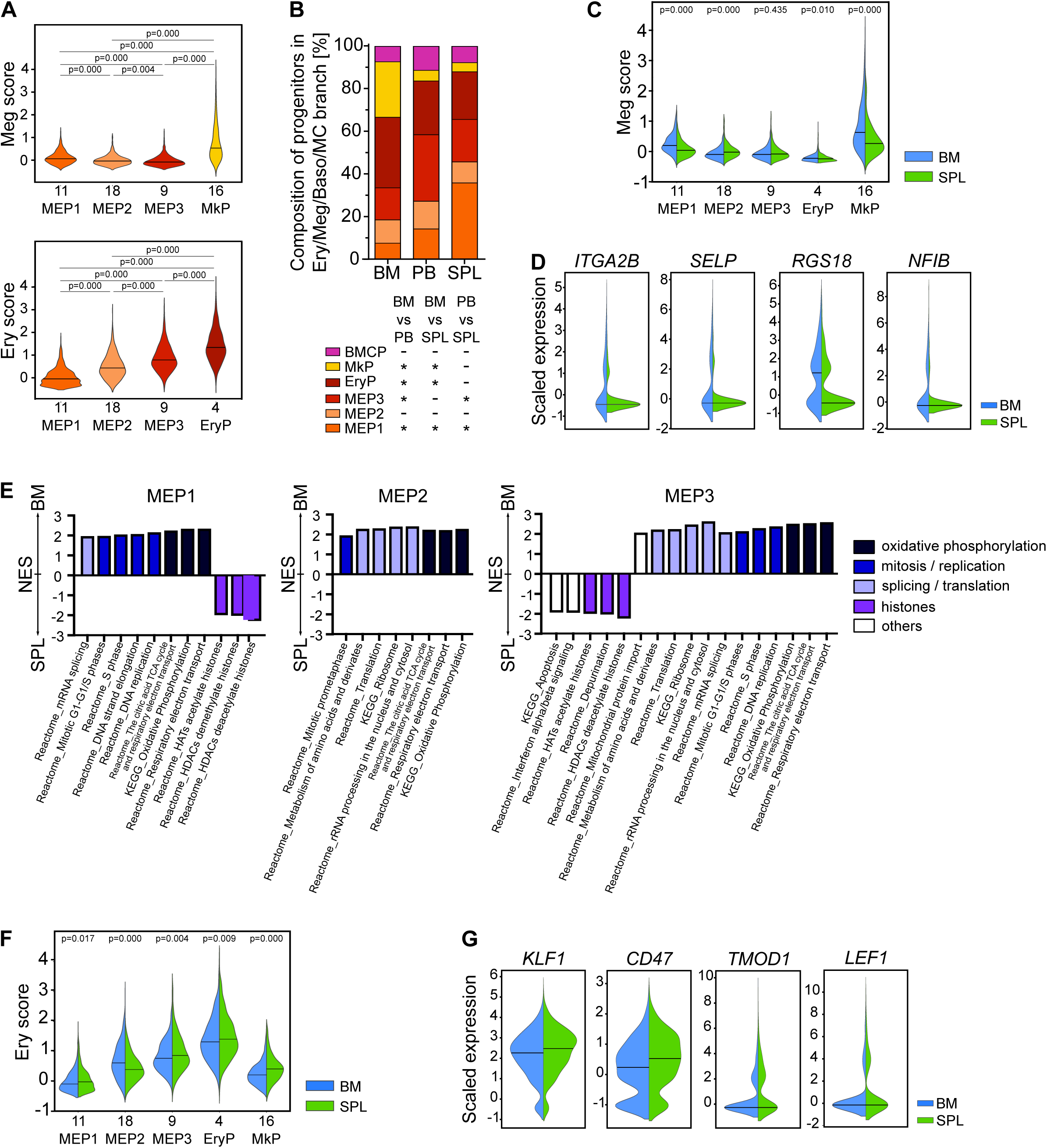
Different balances of Ery/Meg priming in BM and spleen. Analysis of 10x Genomics scRNAseq data from 30,873 cells, combining both donors and all tissues **(A)** or comparing different tissues **(B-G)**. **A)** A score of Meg-lineage priming (‘Meg score’) and Ery-priming (‘Ery score’) was calculated using the average expression of genes known to be up-regulated during early Meg or Ery differentiation. Violin plots show the lineage priming score for each MEP cluster and the latest unipotent progenitor of each lineage. **B)** Bar graph of the relative composition of all clusters within the Ery/Meg/Baso/MC branch (clusters 4, 9, 11, 16, 18, 21) combining datasets of both organs. Data of individual donors is shown in **Figure S3A**. Fisher test. * p<10^-5^. **C)** Split violin plots of the Meg scores for each cluster in the Ery/Meg branch in BM and spleen. **D)** Violin plots of expression of selected Meg-associated genes significantly upregulated in MEP1 cells from BM compared to spleen (FDR<0.05 by EdgeR). **E)** Selected significantly enriched genes sets of MSigDB C2 curated molecular pathways (pre-ranked GSEA, FDR <0.05) in MEP1, MEP2 and MEP3 cells comparing BM and spleen. Gene sets of the same functional category are highlighted by the same colour. Selected gene sets are shown. All gene sets are listed in **Table S8**. **F)** Split violin plots of the Ery scores for each cluster in the Ery/Meg branch in BM and spleen**. G)** Violin plots of expression of selected early erythroid genes significantly upregulated in EryPs from spleen compared to BM (FDR<0.05 by EdgeR). **C,D,F,G:** BM cells (blue) and spleen cells (green); **C,F:** the median in shown as solid line and p-values are indicated as determined by Welch modified t-test. SPL: spleen.

Focusing on Ery differentiation, we found that whereas highly proliferative EryPs were significantly enriched in BM compared to PB and spleen, bipotent MEPs, in particular MEP1, were significantly more abundant in extramedullary tissues (**Figure 3B, S3A**), despite having twice fewer cells actively proliferating (42% S-G_2_-M MEP1 in BM, 21% S-G_2_-M MEP1 in spleen, **Figure 2D**). GSEA analysis confirmed that spleen MEP1/2/3 were negatively enriched for expression of gene sets related to replication, translation and oxidative phosphorylation in line with their reduced cell cycle activity (**Figure 3E, Table S8**). Ery scores were generally higher in spleen than in BM at all stages of differentiation (with the exception of MEP2, **Figure 3F**). Finally, spleen EryPs expressed significantly higher levels of genes associated with early erythroid-biased MEPs, such as *KLF1, CD47, TMOD1 and LEF1* (Psaila et al., 2016) than BM EryP (**Figure 3G, Table S9)**. Taken together, this analysis shows that the balance in Ery/Meg/Baso/MC branch of the spleen is shifted towards an accumulation of slowly proliferating early progenitors, primed towards Ery differentiation. This conformation whereby Ery differentiation in the spleen is partially blocked at steady state but poised to resume in response to stress, is consistent with the role of the spleen in stress erythropoiesis.

### HSC/MPPs in extramedullary sites are transcriptionally poised for activation

Very little is known about the molecular and functional characteristic of HSC/MPPs resident in the human spleen. Analysis by flow cytometry confirmed the presence of adult phenotypic human HSC/MPPs (defined as CD19^-^CD34^+^CD38^-^CD45RA^-^) in all three organs **(Figure 4A)**. Consistent with data in mice (Morita et al., 2011) the frequency of phenotypic HSC/MPPs is 5- to 10-fold lower in PB and spleen than in BM. These cells were highly quiescent residing almost exclusively in G_0_ (**Figure 4B**). Importantly, when transplanting HSPC enriched fractions (Lin-) from spleen into NSG mice, we detected repopulation capacity for at least 14 weeks post-transplantation **(Figure 4C, Table S10)**, demonstrating that although being rare in frequency, the spleen hosts phenotypic HSC/MPPs with repopulation capacity *in vivo*.

**Figure 4.**
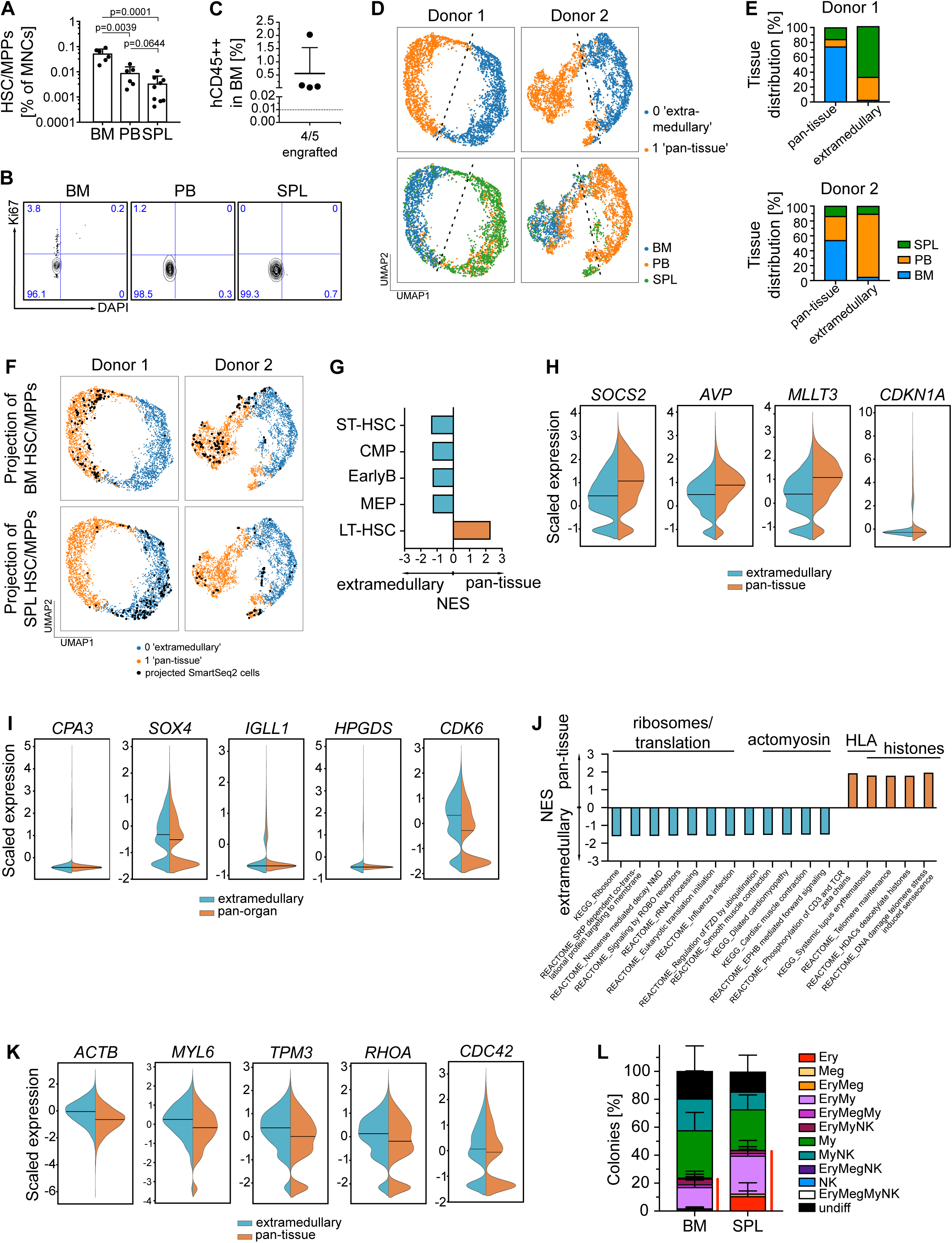
HSC/MPPs from extramedullary sites are in a pre-activated lineage-primed state. **A)** Frequencies of phenotypic HSC/MPPs (CD19-CD34+ CD38-CD45RA-) in mononuclear cells (MNCs) from BM, PB and spleen. Unpaired t-test. **B)** Flow cytometry plots of BM (left), PB (middle) and spleen (right) CD19^-^ CD34^+^ CD38^-^ CD45RA^-^ phenotypic HSC/MPPs in G_0_ (Ki-67^-^DAPI^-^), G_1_ (Ki-67^+^DAPI^-^) and S-G_2_-M (Ki=67^+^DAPI^+^) cell cycle phases. % of cells in each quadrant is indicated. **C)** Lineage-depleted (Lin^-^) cells from spleen were injected intravenously into 5 sublethally irradiated NSG mice. Human engraftment in the BM of recipient mice was analysed 14 weeks after transplantation and defined as positive if percent of human leukocytes (hCD45^++^) in all BM MNCs >0.01. **D-K:** Analysis of 10x Genomics scRNAseq data from 7,563 HSC/MPP Tier 1 cells (donor 1: 3,224 cells, donor 2: 4,339 cells) combining all tissues **D)** UMAPs of HSC/MPP tier1 cells clustered by SAM (kmeans=2; top panels coloured by SAM cluster, bottom panels by tissue). Donors were analysed individually (left panel: donor 1, right panel: donor 2). **E)** Bar graphs of the proportions of BM, PB and spleen-derived HSC/MPP tier1 cells in the pan-tissue and extramedullary SAM clusters. **F)** Projection of Smart-seq2 single-cell transcriptomes of phenotypic HSC/MPPs (same donors and tissues as droplet approach, n=129 from BM donor 1, 165 from spleen donor 1, 242 from BM donor 1, and 74 from spleen donor 2) onto the reference HSC/MPP tier1 subset of cells obtained via 10x Genomics droplet sequencing for the same tissue (only BM or spleen) and donor. Visualisation is shown on same UMAP as in (D). Smart-seq2 cells projections are visualized on UMAP by black dots representing the top projected neighbour of the reference 10x dataset used for the projection. BM (top panel) and spleen (bottom panel). **G-K:** analysis of genes differentially expressed between the pan-tissue (n=3,743 cells) and extramedullary (n=3,820 cells) SAM derived clusters of HSC/MPP tier1 cells. **G)** pre-ranked GSEA of population-specific signatures (LT-HSC and ST-HSC from (Laurenti et al., 2015), other from (Laurenti et al., 2013)) comparing ‘pan-tissue’ and ‘extramedullary’ HSC/MPP tier1 cells. All gene sets enriched (FDR<0.05) are shown. **H-I)** Split violin plots of scaled expression values of selected genes belonging to the LT-HSC **(H)** or ST-HSC **(I)** gene sets enriched in ‘pan-tissue’ HSC/MPP tier1 cells and significantly upregulated in the ‘pan-tissue’ cluster (orange) compared to the ‘extramedullary’ (blue) cluster (FDR<0.05 by EdgeR). **J)** Gene set enrichment of C2 curated pathways (FDR <0.05 by pre-ranked GSEA) comparing ‘pan-tissue’ HSC/MPP tier1 cells with ‘extramedullary’ HSC/MPP tier1 cells. Selected gene sets are show. All gene sets are listed in **Table S13**. **K)** Split violin plots of scaled expression values of selected genes related to actomyosin pathways comparing cells of the ‘extramedullary’ (blue) and ‘pan-tissue’ (orange) clusters. **L)** Single phenotypic HSC/MPPs from BM (n=496 single cells) and spleen (n=234 single cells) were seeded into medium supporting My/Ly/Ery/Meg differentiation. After 3 weeks colonies derived from each single cell were analysed by flow cytometry. Bar graph shows average colony composition for each organ. The red line on the right of each stacked bar indicates all Ery-containing colonies (see also **Figure S4F**). **H,I,K:** in all violin plots the median in shown as solid line. SPL: spleen.

To investigate whether extramedullary HSC/MPPs differ from their BM counterparts, we then analysed how single cells from different tissues distributed in the ‘HSC/MPP Tier1’ space. Tier1 contains the most immature cells in the HSPC landscape as indicated by: i) >85% G_0_/G_1_ phase (**Figure 1F**), ii) highest expression levels of stem cell marker genes such as *MLLT3, SOCS2* and *CDKN1A* (**Figure 1D**) and iii) being the area where most phenotypic HSC/MPPs projected onto **(Table S5)**. We noticed that Louvain cluster 0 was highly overrepresented in spleen/PB and cluster 3 in BM, whereas cells from all organs were found in other Tier1 Louvain clusters (**Figure 2A-B, S2A**). Thus, we re-clustered all cells within Tier1 HSC/MPPs using the ‘Self Assembling Manifolds’ (SAM) algorithm that was designed to identify subtle but biologically relevant differences between subpopulations (Tarashansky et al., 2019). For both donors, this analysis segregated Tier1 HSC/MPPs into two clusters (see methods, **Table S11**). One cluster, which we termed ‘pan-tissue’, contained all BM cells from Tier1 as well as cells from spleen and PB. The other, which we termed ‘extramedullary’ strikingly was exclusively constituted of cells from spleen or PB **(Figure 4D-E)**. Projection of Smart-seq2 data of phenotypic HSC/MPPs confirmed that BM phenotypic HSC/MPPs predominantly project onto the ‘pan-tissue’ cluster, while spleen phenotypic HSCs mainly project on the extramedullary cluster **(Figure 4F)**. Similarly, on combining Tier1 with Tier2 cells, a cluster containing almost exclusively extramedullary cells was identified (data not shown). These data suggest that 2 subtypes of HSC/MPPs coexist in extramedullary tissues: a minority of cells with the molecular features of BM HSC/MPPs and a vast majority with a unique transcriptional phenotype.

To identify the transcriptional differences between ‘pan-tissue’ and ‘extramedullary’ HSC/MPPs, we performed differential gene expression analysis on the droplet-based data of both donors. 761 and 6,599 genes were significantly upregulated (FDR<0.05) respectively in the pan-organ and ‘extramedullary’ cluster (**Table S12**). We detected significant differences in the enrichment of population-specific signatures (Laurenti et al., 2013, 2015) and lineage-associated gene modules (Velten et al., 2017) between both groups (**Figure 4G, Figure S4A, Table S13**). In particular, pan-tissue HSC/MPPs were enriched for a signature of genes associated with long-term HSCs, and correspondingly expressed significantly higher levels of *SOCS2, MLLT3, CDKN1A* **(Figure 4H)** and many histone genes **(Figure S4B)** (Cabezas-Wallscheid et al., 2014; Calvanese et al., 2019; Cheng et al., 2000; Vitali et al., 2015), suggesting that pan-tissue HSC/MPPs contain the most immature quiescent HSCs. In contrast, extramedullary HSC/MPPs were enriched in signatures of genes corresponding to short-term HSCs and lineage-committed progenitors (CMP, EarlyB, MEP, **Figure 4G, Table S13**). Accordingly, *CPA3*, *SOX4*, *IGLL1* and *HPGDS* were all expressed at significantly higher levels in extramedullary HSC/MPPs (**Figure 4I, Table S12)**, indicating a higher degree of lineage-priming than ‘pan-tissue’ HSC/MPPs. Importantly, although being mainly non-dividing (transcriptome based cell cycle assignment, Ki67-DAPI-by flow cytometry), extramedullary HSC/MPPs also showed increased expression of the cell cycle regulator *CDK6* (**Figure 4I, Table S12)**, suggesting their rapid cell cycle entry upon mitogen stimulation (Laurenti et al., 2015; Scheicher et al., 2015). This observation indicates that HSC/MPPs from sites are in a pre-activated state likely enabling faster cell cycle entry upon sensing a mitogenic signal, similar to that of cord blood ST-HSCs (Laurenti et al., 2015). Supporting this hypothesis was the enrichment in extramedullary HSC/MPPs of gene sets associated with ribosomes and translation (**Figure 4J, and Table S13)**. These data indicate that HSC/MPPs are poised for rapid division and lineage differentiation in response to extrinsic stimuli.

HSC/MPPs from the ‘extramedullary’ cluster were further enriched for genes sets involved in the actomyosin cytoskeleton, such as *ACTB*, *ACTG1*, *TPM1/3/4*, *MYL6* and *MYL12A*, suggesting different mechanical properties of HSC/MPPs outside the bone marrow (**Figure 4J and 4K, Table S13**). Other significantly up-regulated genes included the Rho family small GTPases *RHOA*, *RAC1* and *CDC42* (**Figure 4J and 4K)** and their downstream targets *ROCK1* and *2* **(Table S12)**, which are key regulators of actomyosin rearrangements involved in cell migration and cell polarity (Amano et al., 2010). Interestingly, RhoA/Rock I and CDC42 activity is crucial for macrophage chemotaxis (Allen et al., 1998), as well as human and mouse HSPC motility *in vitro* (Fonseca et al., 2010; Yang et al., 2001).

Altogether the transcriptomic data from our droplet-based approach demonstrate that the majority of HSC/MPPs in extramedullary organs i) are poised for division, ii) have upregulated lineage-priming genes and iii) express distinct adhesion and cytoskeletal molecules. Importantly, these findings were validated in our Smart-seq2 sequencing data. Differential expression analysis between single phenotypic HSC/MPPs isolated from BM and from spleen showed similar enrichment of lineage-priming signatures **(Figure S4C, Table S14)** and significantly higher expression of lineage-marker genes such as *CPA3*, *FCER1A*, *ENO1* in spleen (**Table S15**). Several actomyosin genes were also significantly upregulated in spleen compared to BM HSC/MPPs (**Table S15**).

Finally, to functionally verify if the increased lineage priming observed in extramedullary HSC/MPPs affects their differentiation, we performed single cell differentiation assays from single HSC/MPPs (CD19^-^CD34^+^CD38^-^CD45RA^-^ cells) as in (Belluschi et al., 2018a). Interestingly spleen HSC/MPPs produced more colonies containing Ery cells than their BM counterparts (**Figure 4L and S4D-F**). We conclude that consistent with the role of spleen in stress erythropoiesis, spleen HSC/MPPs are molecularly and functionally primed towards Ery differentiation.

## DISCUSSION

The dynamics of extramedullary haematopoiesis in humans has remained understudied due to the difficulty of accessing human samples. Here, using tissues from deceased organ donors, we have mapped the HSPC composition of paired BM, PB and spleen in two individuals at single cell resolution. We report the following key findings. First, distinct balances of HSC/MPP and progenitor differentiation are observed in BM and in extramedullary tissues. Second, the BM is the primary site of progenitor proliferation, in particular for erythro-myeloid lineages. Third, we identify a unique molecular subtype of HSC/MPPs uniquely present in extramedullary tissues, which is quiescent but primed for translation and differentiation. Our findings highlight a poised cellular composition and molecular phenotype of HSPCs in extramedullary organs, sustaining a hierarchical configuration positioned to rapidly contribute to haematopoiesis in response to stress.

The composition of the BM HSPC pool observed here in deceased organ donors is largely similar to that previously reported studies using BM from live donors (Hay et al., 2018; Pellin et al., 2019; Velten et al., 2017). We thus infer that our analysis is overall reflective of steady-state conditions in humans. Nonetheless we cannot exclude that some of the molecular features and/or the generally poised state observed here and discussed below are influenced by cellular responses induced upon death and/or tissue storage (Madissoon et al., 2019).

Our single cell transcriptomic analysis identified distinct structures of the haematopoietic tree in adult haematopoietic tissues. We find that Ly progenitors committed to either B, T or NK cell differentiation were relatively much more abundant in the spleen than in BM. In contrast, dendritic cell progenitors (most likely plasmacytoid DC precursors) and Meg progenitors are almost exclusively found in the BM. The latter likely provides a local source of Meg for efficient platelet production, as platelet release is dependent on the physical interaction of Megs with the BM vascular niche (Grozovsky et al., 2015; Junt et al., 2007).

The existence of multipotent repopulating HSC/MPPs in the human spleen has been postulated from studies in mice (Coppin et al., 2018; Morita et al., 2011; Mumau et al., 2017) and *in vitro* long-term cultures of human splenocytic preparations (Dor et al., 2006). Here we provide evidence that rare but transplantable HSC/MPPs exist in human spleen. In addition, we show that the vast majority of molecularly defined HSC/MPPs possess a distinct transcriptomic phenotype from their BM counterparts. Extramedullary HSC/MPPs have significantly higher levels of Rho/ROCK pathway and actomyosin genes, likely indicating increased motility (Fonseca et al., 2010; Yang et al., 2001) and reflecting interactions with a distinct niche. In addition, their elevated activity of the Rho family GTPase *CDC42* may confer a loss in cell polarity and influence their lineage output and repopulation potential, similarly to aged murine HSCs (Florian et al., 2012).

The spleen is known to contribute extensively to blood regeneration in conditions of stress, such as anaemia or injury-induced stress erythropoiesis, or in response to infections, when HSPCs from the BM actively migrate into the spleen and proliferate there (Griseri et al., 2012; Harandi et al., 2010; Lenox et al., 2005; Masuya et al., 2014; O’Malley et al., 2005). The role of tissue-resident HSPCs in these responses has remained elusive. Our data altogether indicate that not only does the spleen provide a niche for BM cells during emergency haematopoiesis, but that it also contains molecularly distinct resident HSPCs which are poised to efficiently divide in response to stress. This is supported by the fact that unilineage progenitors of all branches are not actively dividing in the spleen. In addition, HSC/MPPs uniquely present in extramedullary tissues are almost exclusively non-dividing but express higher levels of ribosomes, translational machinery genes and *CDK6*, the hallmarks of pre-activated cells primed to divide fast in response to a stimulus (Laurenti et al., 2015; van Velthoven and Rando, 2019). Interestingly, mouse splenic HSCs mostly reside in G_1_ (Coppin et al., 2018) and cycle twice as often as BM HSCs (Morita et al., 2011), confirming distinct cell cycle properties of BM and extramedullary HSCs across species. Consistently with the role of the spleen in stress erythropoiesis (Harandi et al., 2010; Inra et al., 2015; Lenox et al., 2005), we observe that the cellular and molecular balance in the Ery/Meg/Baso/MC branch in the spleen is tipped towards the earliest Ery progenitors, which accumulate and cycle significantly less than in BM. We speculate that these cells may be a stress-responsive progenitor population, and future experiments will have to address if they are akin to the splenic stress BFU-E described in mice (Harandi et al., 2010; Perry et al., 2007). In addition, spleen HSC/MPPs were functionally primed to produce more Ery cells *in vitro*, indicating that this population may also contribute to stress erythropoiesis.

Collectively our data highlight that HSPC hierarchies differ in distinct anatomical locations. It is well known that the BM harbours and provides a protective environment for the most quiescent HSCs, which are thought to contribute to blood production almost exclusively during extreme stress situations (Foudi et al., 2009; Nakamura-Ishizu et al., 2014; Wilson et al., 2008). From our data, we speculate that the spleen is a complementary reservoir of quiescent HSC/MPPs and slowly-proliferating progenitors, that is naturally positioned at steady-state for fast production of differentiated progeny in response to milder stresses. Future studies will address whether the particular configuration of the extramedullary HSPC hierarchy is maintained by the absence of proliferative signals or by the splenic niche actively inhibiting differentiation. Overall, our data represent a comprehensive resource to understand the molecular underpinning of extramedullary haematopoiesis and its contribution to a number of diseases.

## ACKNOWLEDGEMENTS

We would like to thank the deceased donors, their families and the Cambridge Biorepository for Translational Medicine for access to human tissue (BM, PB and spleen samples), the Cambridge NIHR BRC Cell Phenotyping Hub for their flow cytometry services and advice, and the CRUK Cambridge Institute genomics centre for sequencing; E.L. is supported by a Sir Henry Dale fellowship from Wellcome/Royal Society. Research in E.L.’s laboratory is supported by Wellcome, BBSRC, EHA and Royal Society. Research in E.L. and B.G. laboratories is supported by core support grants by Wellcome and MRC to the Wellcome-MRC Cambridge Stem Cell Institute. Research in H.T.’s laboratory is supported by KAKEN Grant-in-Aid for Scientific Research for Young Scientists (A) (15H05669) and challenging Exploratory Research (18K19520). N.M. is supported by a DFG Research Fellowship (ME 5209/1-1), NKW and BG were supported by grants from Bloodwise, Wellcome, CRUK and MRC. A.C. is supported by Wellcome, A.S. by a Cambridge Cancer Centre fellowship and K.T.M by the Chan Zuckerberg Initiative.

## CONTRIBUTIONS

Conceptualization: N.M. and E.L.; Methodology: N.M., A.S., N.W. A.C. and K.M.; Formal analysis: N.M., H.B. K.S. and E.L.; Investigation: N.M., A.S., N.W., A.C and K.M.; Resources: K.S.P.; Writing – Original Draft: N.M, H.B. and E.L.; Writing – Review & Editing: H.T., N.M, E.L.; Visualization: N.M., H.B. and E.L.; Supervision: H.T., K.S.P., B.G., and E.L.: Project administration: E.L.; Funding acquisition: B.G. and E.L.

## MATERIALS AND METHODS

### Human samples

Steady-state bone marrow (BM), spleen and peripheral blood (PB) were taken from consented deceased healthy organ donors **(see Table S1)** by the Cambridge Biorepository for Translational Medicine based in the Department of Surgery at the Cambridge University Hospitals NHS Trust Addenbrooke’s Hospital in Cambridge. Organ donation occurred either after circulatory death (DCD) or after brain stem death (DBD). Tissue sampling was in accordance with regulated procedures approved by the relevant Research and Ethics Committees (REC 15/EE/0152) and was obtained after informed consent from the donor family. For cell cycle staining by flow cytometry peripheral blood from a healthy living blood donor was used. Blood was collected with informed consent at the NHS Blood and Transfusion (NHSBT) Centre in Cambridge in accordance with regulated procedures approved by the relevant Research and Ethics Committees (REC 12/EE/0040).

### Sample preparation

For isolation of mononuclear cells (MNCs) from BM and PB, samples were diluted 1:1 in PBS prior to density gradient centrifugation on Pancoll (PAN-Biotech). The MNC fraction was collected and red blood cells were lysed in RBC lysis buffer (BioLegend) for 15 min at 4 °C. After splenic tissue collection, samples were immediately placed into 4°C UW solution and kept at 4°C until processing (within 12 hours of collection). Single-cell suspensions were obtained by placing small sections of spleen (<5mm^3^) with cold phosphate buffer into gentleMACS™ C tubes and utilizing a benchtop gentleMACS™ dissociator (Miltenyi Biotec). MNCs were enriched through gradient separation (density gradient: 1.077 ± 0.001 g/ml, STEMCELL Technologies) followed by red cell lysis. MNC cell suspensions were frozen as aliquots of up to 5×10^7^ viable cells per vial in FBS/10% DMSO and stored at −150 °C until use.

### Flow cytometry

MNCs were thawed by drop wise addition of pre-warmed IMDM (Life Technologies) + 0.1 mg/ml DNase (Worthington) + 50% Fetal Bovine Serum (FBS, Life Technologies). If required, magnetic CD34^+^ selection was performed prior to antibody staining, using the CD34 MicroBead Kit (Milentyi Biotec) and manual separation on LS columns (Milentyi Biotec). Cells were stained for 20 min at RT in the dark using antibodies diluted in PBS + 3% FBS as listed in **Table S16**. Zombie Aqua or Zombie UV (Biolegend) was used as viability dye. Prior to flow cytometry, samples were washed twice in PBS + 3% FBS, then resuspended in PBS + 3% FBS and filtered through a 20 μm cell strainer.

Cell cycle analysis by flow cytometry was performed on BM and spleens from deceased organ donors, while peripheral blood was obtained from healthy living blood donors. Cell surface markers were stained as described above, followed by fixation in Cytofix/Cytoperm (BD) for 15 min on ice according to manufacturer’s protocol. After two washing steps in Perm/Wash (BD), intranuclear staining was performed at 4 °C overnight in the dark using the FITC anti–human Ki-67 antibody (BD) diluted in Perm/Wash. The next day, DNA content was stained by 20 min incubation in 0.5 μg/ml DAPI (BioLegend) in Perm/Wash at RT, followed by one wash with Perm/Wash. Samples were resuspended in PBS + 3% FBS and analysed by flow cytometry.

Samples were sorted using the BD FACSAria Fusion and FACSAria III sorters at the NIHR Cambridge BRC Cell Phenotyping Hub. Single-cell sorts were performed using single-cell purity and index-sorting modes, while purity mode was chosen for bulk sorts. For flow cytometry analysis the BD LSRII and LSRFortessa cytometers were used. High-throughput analysis was performed using the BD High Throughput Sampler (HTS). Data were analysed using FlowJo software v9.9 or v10 (Tree Star).

### Antibody panels

Clone, supplier and catalogue number for each antibody can be found in **Table S16**. Antibody panels A-D were used to sort and phenotype subsets of hematopoietic stem and progenitor cells. **Panel (A):** CD19/BV785, CD34/APC-Cy7, CD38/PE-Cy7, CD90/APC, CD45RA/A700, CD45f/PE-C5, CD10/BV421, CD71/FITC, CD117/BV650, CLEC9A/PE, Zombie Aqua; **Panel (B):** CD19/BV785, CD34/APC-Cy7, CD38/PE-Cy7, CD90/APC, CD45RA/A700, CD45f/PE-C5, CD41a/BV510, CD71/FITC, CD117/BV650, CLEC9A/PE, Zombie UV; **Panel (C):** CD45RA/FITC, CD90/PE or CD90/APC, CD45f/PE-Cy5, CD38/PE-Cy7, CD10/APC or CD10/BV421, CD19/A700, CD34/APC-Cy7, CD7/BV421, Zombie Aqua; **Panel (D):** CD19/BV785, CD34/APC-Cy7, CD38/PE-Cy7, CD36/APC, CD45RA/A700, CD71/PerCP-Cy5.5, GlyA/BV421, CD90/FITC, CD45/BV605, CD135/PE, Zombie Aqua. For cell cycle analysis, **panel (E)** was used: CD45RA/PE, CD90/APC, CD45f/PE-Cy5, CD38/PE-Cy7, CD10/BV421, CD19/A700, CD34/APC-Cy7, CD7/BV421, KI-67/FITC, DAPI. After exclusion of dead cells based on Zombie dyes, HSPCs populations were defined as previously described: total HSPC pool (CD19-CD34+), phenotypic HSC/MPPs (CD19^−^CD34^+^ CD38^−^CD45RA^−^). Colonies from My, Ly, Ery and Meg single-cell differentiation assays were stained with **Panel (F):** CD45/PE-Cy5, CD41/FITC, CD11b/APC-Cy7, CD14/PE-Cy7, CD15/BV421, GlyA/PE, CD56/APC. Prior to transplantation into mice, efficiency of lineage depletion was analysed using **Panel (G):** CD3/FITC, CD5/PE, CD45f/PE-Cy5, CD34/APC-Cy7, CD38/PE-Cy7, CD90/APC, CD45RA/A700, Streptavidin/BV510 (BioLegend, dilution 1:400), Zombie UV. At time point of final analysis, mouse bone marrow was stained with antibody **panel (H)**: GlyA/PE, CD19/FITC, CD45/PE-Cy5, CD14/PE-Cy7, CD33/APC, CD19/A700, CD45/BV510, CD3/APC-Cy7.

### RNA-sequencing by Smart-seq2 protocol

In parallel to scRNA-seq of CD34+ HSPCs using the 10x Genomics platform, single-cell sorted phenotypic HSC/MPPs from the same donors were sequenced with the Smart-seq2 protocol. For donor 1 only BM and spleen phenotypic HSC/MPPs were sequenced using this method, for donor 2 phenotypic HSC/MPPs from all three tissues were sequenced using Smart-seq2 (**see Table S2).** Library preparation of single cells was adapted from the Smart-seq2 protocol of Picelli et al (Picelli et al., 2014). First, individual cells were sorted into 4 μl of lysis buffer per well of 96-well PCR plates. Lysis buffer contained 0.22% (v/v) Triton X-100, 11.5 U/μl RNase inhibitor (Clontech), 5 mM DTT, 1 mM dNTP in nuclease-free water (Ambion, Life Technologies). During the sort, the information on cell surface marker levels was recorded as index .csv file for each cell. Plates were spun down immediately after sort and stored at −80 °C until library preparation. After addition of 10 μM OligodTs (Sigma) and ERCCs at a final concentration of 1:30,000,000 (Ambion, Life Technologies), annealing was performed for 3 min at 72 °C. Next, reverse transcription was carried out using 50 U/well SmartScribe (SmS) RT enzyme (Takara) + 1x SmS First Strand buffer (Takara) + 5 μM TSO (Qiagen). Amplification of cDNA (22 amplification cycles) was performed using 0.55 μM ISPCR primers (Sigma) and 1 x KAPA HiFi Hotstart mix (Roche). After purification of the PCR product with a 1: 0.6 ratio of PCR product to AMPure XP beads (Beckman Coulter), the success of cDNA preparation was confirmed using a 2100 Bioanalyzer with High-sensitivity DNA chip (Agilent). Illumina library preparation with the Nextera XT DNA Library Prep Kit (Illumina) was carried out following manufacturer’s instructions. Size distribution of the libraries was analyzed on a High-sensitivity DNA chip (Agilent) and the concentration of indexed libraries was quantified following the protocol of the KAPA library quantification kit (Kapa Biosystems). For each donor libraries from all tissues were combined prior to sequencing and sequenced on the Illumina Hiseq 4000 system in 2 lanes.

Resulting reads from each donor were independently aligned and mapped against Ensembl genes (release 81) (Zerbino et al., 2018) using GSNAP (version 2015-09-29) (Wu and Nacu, 2010) and quantified using HTSeq (version 0.6.0) (Anders et al., 2015). Quality control filters were applied rejecting cells with less than 50,000 (donor 1) or 200,000 (donor 2) reads mapping to nuclear genes which in turn had to be at least 10% of the mapping reads in the cell. Additionally, the maximum allowed fraction of cells mapping to mitochondrial genes was set at 20%. The levels of technical variance were estimated using the ERCC spike-ins as described by Brennecke et al. (2013) (Brennecke et al., 2013) with highly variable genes (HVGs) being defined as having the squared coefficient of variation exceeding technical noise. The raw sequencing reads and gene count tables were deposited at the National Center for Biotechnology Information GEO (accession number GSE143567).

### Combined donor Smart-seq2 data analysis

Prior to differential gene expression analysis, the Smart-seq2 data filtered count matrices from both donors were combined and HVGs determined using the method described by Brennecke et al. (2013). Principal component analysis (PCA) was calculated on the HVGs and the cells falling outside the upper and lower limits of the boxplot for the three first principle components were removed as outliers. Differential expression tests were performed using the R packages DESeq2 (Love et al., 2014) with donor identity preserved as a covariate in the design matrix. The batch correction method Combat, was then applied as described in Leek et al. (2012) (Leek et al., 2012) after which the HVG set was recomputed, using the method described in Satija et al. (2015) (Satija et al., 2015). This recomputed HVG set was then subsequently used to recompute PCA and the first two components plotted.

### RNA-sequencing using 10x Genomics platform

For each tissue and donor 7516 – 17500 CD34+ HSPCs **(see Table S2)** were sorted into 300 ul PBS + 3%FCS and kept on ice until library prep. After cells were spun down, they were resuspended in 47 ul PBS + 0.04% BSA and further processed for single-cell sequencing using the Chromium™ Single Cell 3’ Library & Gel Bead Kit v2 (10x Genomics) following the manufacturer’s protocol. For each donor libraries from all three tissues were combined prior to sequencing and sequenced on the Illumina Hiseq 4000 system in 2 (donor 1) or 3 lanes (donor 2). The resulting scRNA-seq reads were aligned to the GRCh37 (hg19) reference genome and quantified using cellranger count (CellRanger pipeline version 2.1.1), giving an average cell recovery of 33-38% for each donor and tissue **(Table S2)**. Multiplets were estimated using the scrublet Python package (Wolock et al., 2019) and subsequently removed.

### Data filtering, normalization and scaling

In the 10x datasets, for both donors, genes being expressed in fewer than 3 cells and cells expressing fewer than 500 genes or cells with 10% or more of its expressing genes being mitochondrial were excluded. Filtered data was normalized (1e4 counts per cell), logarithmized, scaled and the HVGs determined using the Seurat method (Satija et al., 2015). These steps were applied independently to the 10x count matrix of each individual donor, through the use of the respective methods as implemented in the Scanpy Python package (Wolf et al., 2018). Datasets submitted to the pSmart above described are hereby referred to as ‘scaled’.

### Seurat alignment

The 10x count matrices from both donors (after multiplet filtering) were processed with the Seurat (version 2.3.4) R package (Stuart et al., 2019). Each dataset was independently normalized, scaled and had its HVGs determined (Satija et al., 2015). Thereafter, the union of the top 3,000 HVGs from each set was used to perform a canonical correlation analysis. The top 20 canonical correlation vectors were chosen to be used in the method to align of the cells from both datasets. Following the alignment, the Louvain algorithm for community detection (Blondel et al., 2008) was then used to partition the connectivity graph into cell groups and the dimensionality technique t-distributed Stochastic Neighbor Embedding (t-SNE) (Maaten and Hinton, 2008) was computed for the aligned datasets. The resulting combined and processed dataset was exported as an anndata object for further processing with the Scanpy (Wolf et al., 2018) Python package and hence referred to as ‘Seurat-aligned dataset’. Cell communities (henceforth clusters) were defined via the Louvain algorithm using the resolution parameter at 1.77.

### Scanorama integration

The Python package Scanorama (Hie et al., 2019) was also used to integrate the data from both donors. For this purpose the independently scaled 10x datasets, as previously described, were subset to the intersection of the HVGs across the two datasets (3572 genes). These individual donor subsets were used as input for the method and yielded an integrated dataset, henceforth referred to as, ‘Scanorama-aligned dataset’ which was subsequently processed further downstream. Louvain clustering was performed as above with the ‘Scanorama-aligned dataset’, setting the resolution parameter at 1.5 to yield 26 clusters.

### Dimensionality reduction and visualization

Both the individual donor scaled datasets and the combined (Seurat-aligned and Scanorama-aligned) datasets were submitted to dimensionality reduction techniques. These were PCA, diffusion maps (Coifman and Lafon, 2006) and uniform manifold approximation and projection (UMAP) (McInnes et al., 2018). In addition, the ForceAtlas2 force-directed graph (FDG) algorithm (Jacomy et al., 2014) was applied to the k-nearest neighbour graph of the datasets to also provide further embeddings.

### Differential expression

Differential expression tests were performed using the R packages DESeq2 (Love et al., 2014) for Smart-seq2 sequencing data and edgeR (Robinson et al., 2010) using the likelihood ratio test for the 10x sequencing data. Additionally, the Wilcoxon rank sum (WRS) test, as implemented in Scanpy, was used to test differential expression between groups of cells.

### Clustering and lineage characterization

Characterization of the clusters defined via the Louvain algorithm was achieved through i) rank gene groups WRS tests using a one-versus-rest strategy, thus retrieving the most differentially expressed genes in a cluster relative to the remainder clusters; ii) the expression of known gene markers for specific lineages within the Louvain clusters was further investigated by plotting them onto the previously generated embeddings and verifying their co-localization with respect to the defined clusters.

### Pseudotime and Cell Cycle scoring

Pseudotime was calculated from the diffusion map of the combined donors dataset (excluding clusters 23, 24, 25) using diffusion pseudotime (DPT) (Haghverdi et al., 2016). The root cell was selected from the BM cells on cluster 3 (HSC/MPP Tier1). The scores obtained from this method were converted into a rank for better visualization when overlaid in an embedding. The Scanpy implementation of the score genes cell cycle method presented by Satija et al, 2015 (Satija et al., 2015) was used to calculate S phase and G_2_-M phase scores.

### Cell projections

The cells from the Smart-seq2 experiment were projected onto the cells from the 10x experiment (here dubbed reference set) using scmap (Kiselev et al., 2018). Consequently, the datasets were split by donor and tissue and their respective raw count tables were processed as input. Projections were calculated for each respective donor-tissue pair. Visualizations were produced by highlighting the top neighbour cell in an annotated embedding of the reference for each Smart-seq2 projecting cell. Additionally, for each projecting cell the most frequent cluster annotation within the nearest 15 cell projections in the reference was transferred back to that Smart-seq2 cell.

### Self-assembly manifolds of Tier 1 cells

The subset of cells here defined as Tier 1 (louvain clusters 0, 3, 10, 14, 19, based on >85% cells in G_0_-G_1_ by cell cycle assignment as above) was submitted to the Self-assembling manifold (SAM) approach (Tarashansky et al., 2019) with the aim of getting better resolution in cell segregation. SAM was then independently applied to the Tier 1 cell subset of the 10x raw counts from each donor and UMAP embeddings were generated. K-means (k=2) clustering was subsequently computed after visual inspection of the UMAP plots. Additionally, cell projections from the Smart-seq2 cells on these Tier 1 cell subsets were also computed and visualized as described above.

### Gene set enrichment analysis

Gene set enrichment analysis (GSEA) was performed with the GSEA software (v3.0) using pre-ranked gene lists. The ranked lists were derived from differential expression results and based either on the Wald statistic (from DESeq2 results of Smart-seq2 data) or log(pvalue) multiplied by 1 if log2FC>0 and by −1 if log2FC<0 (for edgeR results from 10x data). Enrichment was tested for C2 curated genesets of the MSigDB database v7.0 (Subramanian et al., 2005), published population-specific signatures (Laurenti et al., 2013, 2015) and lineage-priming modules (Velten et al., 2017) as reference gene sets.

### My, Ly, Ery and Meg single-cell differentiation assay

Single-cell assays testing for My, Ery, Meg and Ly differentiation were performed as described in (Belluschi et al., 2018b). One day before the sort, MS5 cells at passage 10-13 (Itoh et al., 1989) (imported from Prof Katsuhiko Itoh at Kyoto University) were seeded at 3000 cells per well of a 96-well flat-bottom plate (Nunc) in 100 μl Myelocult H5100 medium (Stem Cell Technologies) + 1% Pen/Strep (Life technologies). The next day, medium was replaced by 100 μl/well StemPro-34 SFM medium (Life Technologies) supplemented with 3.5 % StemPro-34 nutrients (Life Technologies), SCF 100 ng/ml, Flt3-L 20 ng/ml, TPO 100 ng/ml, IL-6 50 ng/ml, IL-3 10 ng/ml, IL-11 50 ng/ml, GM-CSF 20 ng/ml, IL-2 10 ng/ml, IL-7 20 ng/ml; all Miltenyi Biotec), erythropoietin (EPO) 3 units/ml (Eprex, Janssen-Cilag), h-LDL 50 ng/ml (Stem Cell Technologies), 1% L-Glutamine (Life Technologies) and 1% Pen/Strep (Life Technologies). Single phenotypic HSC/MPPs were index-sorted into each well and incubated at 37 °C for 3 weeks prior to analysis by flow cytometry. If necessary, medium was topped up with 50 μl StemPro medium plus supplements at the half time of culture. After 3 weeks in culture, single-cell derived colonies were transferred through a 30-40 μm plate filter (Pall Laboratory) into 96 U-bottom plates, to exclude co-cultured MS5 cells. Cells were next stained with antibody panel (F) for 20 min at RT in the dark, washed once with PBS + 3% FBS and resuspended in 100 μl/well of PBS + 3% FBS for high-throughput flow cytometry analysis as previously described in (Belluschi et al., 2018b). A single cell was defined as giving rise to a colony if the sum of cells detected in the CD45^+^ and GlyA^+^ gates was > 30 cells. Individual colonies were identified as positive if they contained at least the indicated number of surface-marker positive cells: Ery (CD45^-^GlyA^+^) > 30 cells, Meg CD41^+^ > 20 cells, My colonies (sum of CD45^+^CD14^+^ and CD45^+^CD15^+^) > 30 cells, NK colonies (CD45^+^CD56^+^) > 30 cells. Colonies were identified as undifferentiated (undiff) if the sum of cells in CD45^+^ and GlyA^+^ gates exceeded 30, but the cells could not be identified as positive for any lineage (Ery, Meg, My, NK) using the criteria above. For visualization purposes, EryMegMy colonies are the sum of EryMegMy and MegMy colonies, the latter of which only appeared at proportions <1% in two of the samples. For the same reason the low fraction of EryNk (<1% of all colonies in one sample only) were combined with EryMegNK colonies and are labeld as ‘EryMegNK’.

### Mice

Mice of the NOD.Cg-PrkdcscidIl2rgtm1Wjl/SzJ strain (NSG) were obtained from Charles River or bred in-house. All animals were housed in a Specific-Pathogen-Free (SPF) animal facility and all experiments were conducted under the regulations of the UK Home Office. This research has been regulated under the Animals (Scientific Procedures) Act 1986 Amendment Regulations 2012 following ethical review by the University of Cambridge Animal Welfare and Ethical Review Body (AWERB).

### Xenotransplantation assays

Xenotransplantation was performed on age-matched female NSG mice (age of 7–14 weeks at time of transplantation, details see **Table S10**). One day before transplantation mice were pre-conditioned by sublethal irradiation (2.4 Gy). On the day of transplantation spleen MNCs were thawed and lineage (CD2, CD3, CD11b, CD14, CD15, CD16, CD19, CD56, CD123, and CD235a) positive cells were depleted using the Lineage Cell Depletion Kit (Milentyi Biotec) and magnetic separation via LS columns (Milentyi Biotec). Efficiency of lineage depletion was verified by flow cytometry using staining panel G and was 94±3 % for all samples.

1.5 x10^6^ – 2.5 x10^6^ of lineage-depleted (Lin-) cells were resuspended in 250 ul PBS + 0.1% Pen/Strep (Life Technologies) and intravenously injected into the tail vein of recipient mice.

Mice were analyzed at 14 weeks post transplantation. For this purpose, both tibias and femurs were harvested, bone marrow was flushed with IMDM + 5% FBS and cells were stained with antibody panel H.

To allow confident detection of human leukocyte engraftment, we stained all samples with two distinct antibodies against CD45. Cells were only considered human leukocytes if they were positive for both CD45 antibodies (CD45^++^) as previously described in (Belluschi et al., 2018b). Threshold of engraftment was set to ≥ 0.01 % CD45^++^ cells with at least 30 cells recorded in the CD45^++^ gate. Cells were counted as My lineage if CD45^++^CD33^+^ ≥ 20 cells; Ly lineage if CD45^++^CD19^++^ ≥ 20 cells (positive for two distinct CD19 antibodies); Ery lineage: CD45^−^GlyA^+^ ≥ 20 cells.

### Sample size determination and study design

In all experiments the maximal possible sample size was used, considering limited the availability of primary human samples and rarity of the HSPC populations. Single-cell differentiation experiments were repeated at least three times with independent biological samples. At least 40 single cells were seeded per condition and biological replicate to ensure sufficient power of the assays.

### Statistical analysis

After examination of data distribution and variance between groups, the appropriate statistical tests were used. Statistics on the comparison of cluster composition between tissues was performed using a Fisher’s exact test of the combined numbers of cells from both donors in each tissue and cluster/group of clusters. Statistics on HSC/MPP frequencies across tissues as determined by flow cytometry was done using a two-tailed unpaired t-test. Additional statistics for each bioinformatic analysis are described in each relevant section.

### Data availability and accession

Raw sequencing files and associated metadata for 10x Genomics experiments were deposited at the European Nucleotide Archive. These are accessible via BioStudies with the accession identifier SUBS4: https://www.ebi.ac.uk/biostudies/studies/S-SUBS4.

Raw sequencing files, HTSeq counts and associated metadata for the Smart-Seq2 dataset have been deposited at GEO with the accession number <u>GSE143567</u>.

