## Supplemental Data for "Quantitative and molecular differences distinguish adult human medullary and extramedullary haematopoietic stem and progenitor cell landscapes"

*equal contribution

This file includes:

Supplementary Figures S1 to S4

Legends to Supplementary Tables S1 to S16

Supplementary references

### SUPPLEMENTARY FIGURES

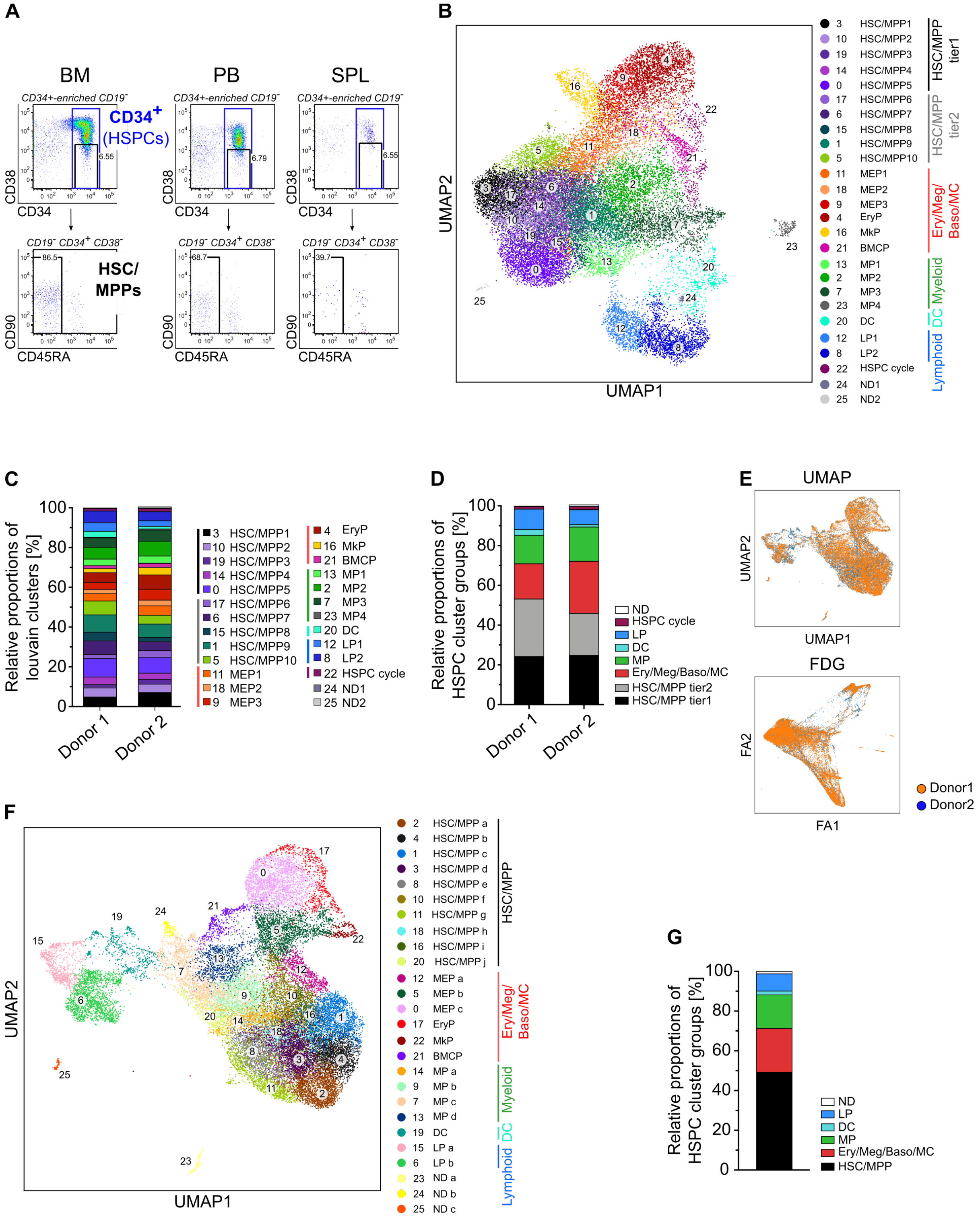


**Figure S1. The adult human HSPC landscape across BM, PB and spleen: related to Figure 1. A)** Flow cytometry plots of BM, PB and spleen samples from donor 1 show the gates used to isolate CD19$^-$ CD34$^+$ HSPCs and CD19$^-$ CD34$^+$ CD38$^-$ CD45RA$^-$ phenotypic HSC/MPPs. **B)** UMAP of the HSPC landscape of all 30,873 cells sequenced with 10x

Genomics single cell 3’ RNAseq, combining all tissues and both donors. Colours and numbers indicate clusters of transcriptionally distinct cells as determined by Louvain clustering. Clusters were annotated using known lineage and stem cell marker genes found amongst the most differentially expressed genes in each cluster (**Table S4**). In addition, clusters were grouped into 6 main HSPC groups as indicated on the right: HSC/MPP tier1, HSC/MPP tier2, Ery/Meg/Baso/MC, Myeloid progenitors, DC progenitors and lymphoid progenitors. ND: clusters which identity could not be defined using known marker genes. HSC/MPP: haematopoietic stem cell/multipotent progenitor; MEP: Megakaryocyte-erythroid progenitor; EryP: erythroid progenitor; MkP: megakaryocytic progenitor; BMCP: Basophil/mast cell progenitor; MP: myeloid progenitor; DC: dendritic cell progenitor; LP: lymphoid progenitor. **C)** Relative proportion of each cluster within the HSPC landscape, combining all three organs, for donor 1 (left) and donor 2 (right). **D)** Cluster composition using the groups defined in (B) for each donor. HSPCs from all organs are combined. **E-G)** Scanorama was used as an alternative method to Seurat alignment to integrate the 10x datasets of all tissues from both donors. **E)** UMAP (top plot) and FDG (bottom plot) of the HSPC landscape after integration with Scanorama. Cells from each donor are depicted by individual colours. **F)** Annotated UMAP of the HSPC landscape after integration with Scanorama. Colours and numbers indicate clusters of transcriptionally distinct cells as partitioned by Louvain clustering. Clusters were annotated as in B). **G)** Cluster composition after cell alignment of all tissues and both donors with Scanorama using the groups as defined in (C). Overall Scanorama resulted in similar clusters and cluster compositions as Seurat **(Figure 1).**

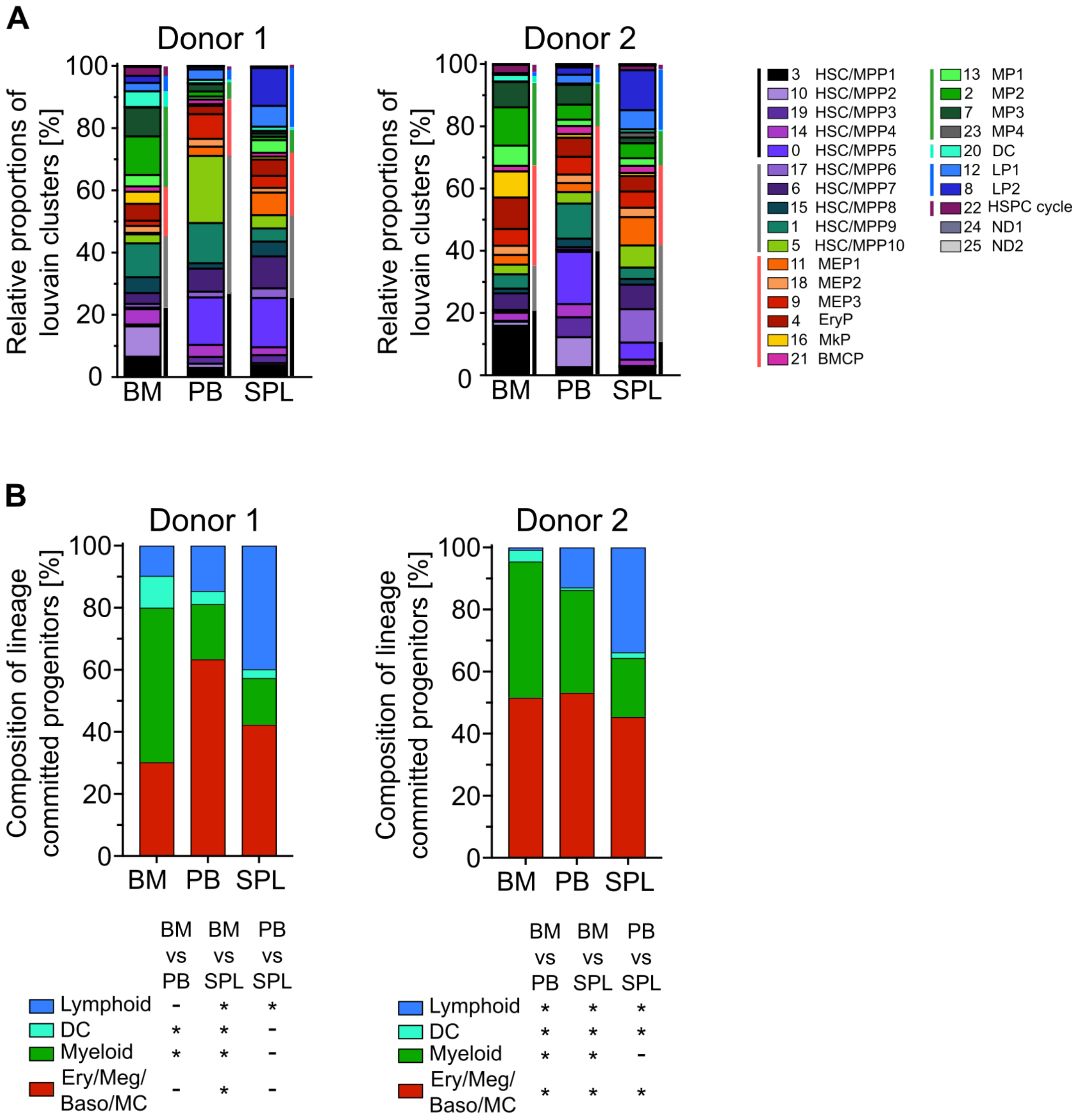


**Figure S2. Similar changes in BM, PB and spleen HSPC cluster composition in both donors: related to Figure 2. A)** Relative cluster composition in BM, PB and spleen. Data from donor 1 is shown on the left, and data from donor 2 on the right. **B)** Bar graph of the relative composition of lineage-committed progenitors in BM, PB and spleen, in each individual donor. Each branch was defined as shown in Figure 1B. Fisher test. * $p<10^{-5}$. SPL: spleen.

**A**

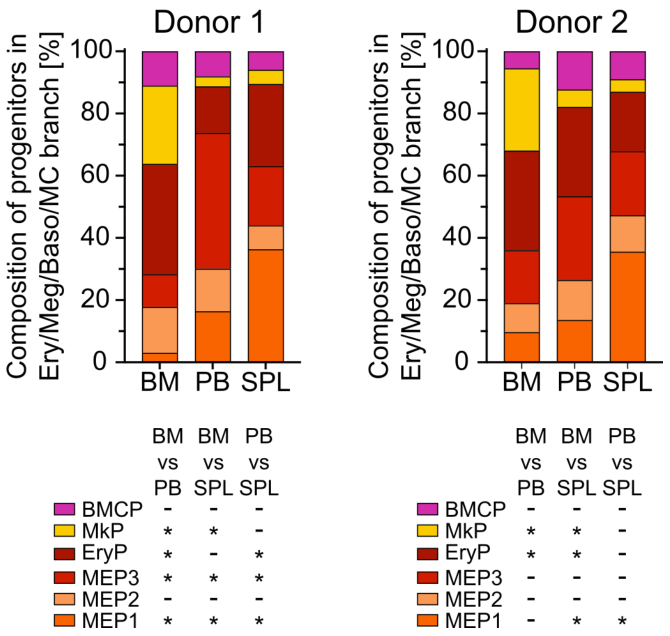


**Figure S3. The composition of the Ery/Meg/Baso/MC branch differs across tissues: related to Figure 3. A)** Bar graphs of the relative composition of all clusters within the Ery/Meg/Baso/MC branch (cluster 4, 9, 11, 16, 18, 21) for each organ. Fisher test. * $p<10^{-5}$. SPL: spleen.

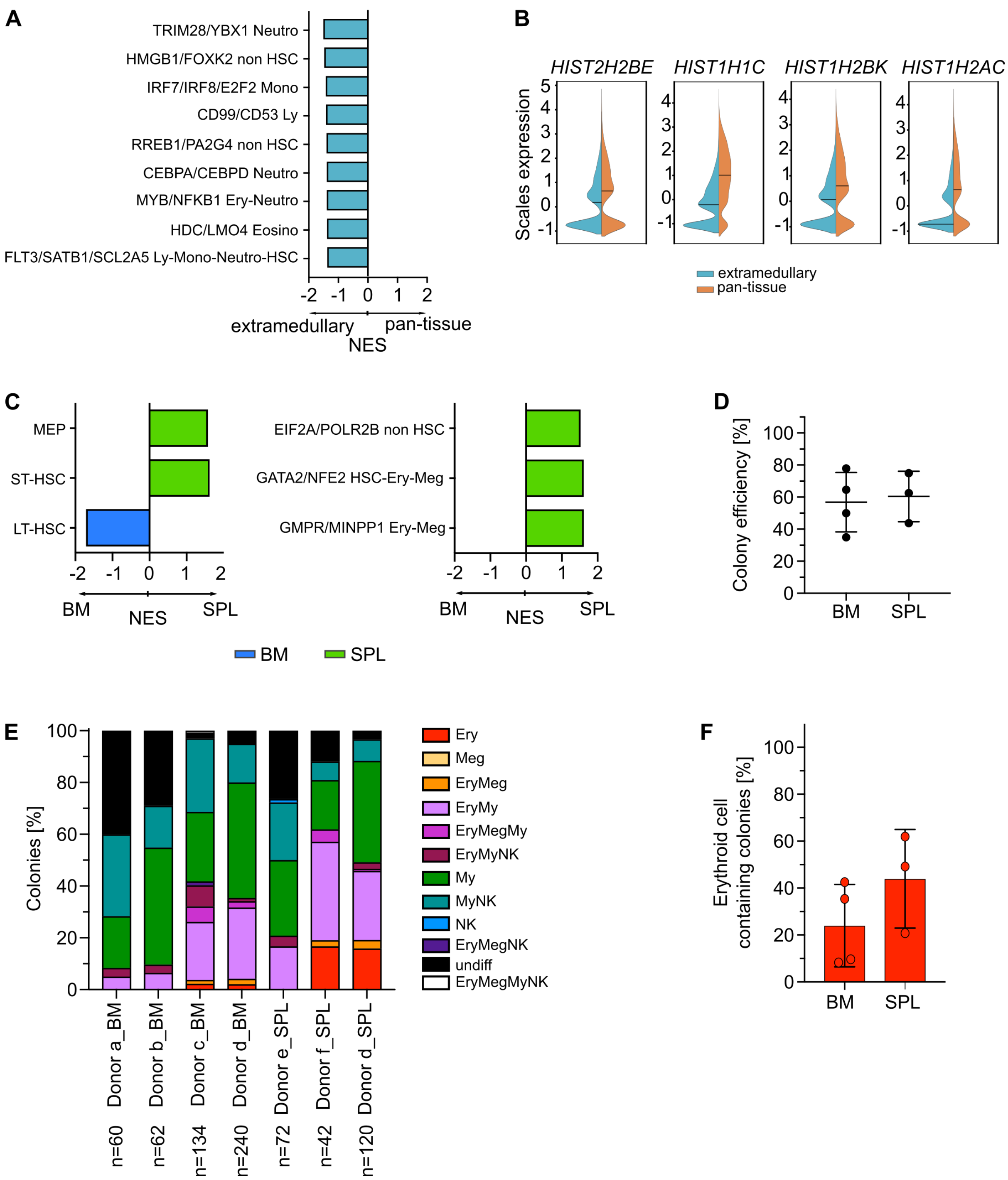


**Figure S4. HSC/MPPs from different haematopoietic tissues are transcriptionally and functionally diverse: related to Figure 4. A)** Pre-ranked GSEA of lineage-priming modules (Velten et al., 2017) comparing 'pan-tissue' HSC/MPP tier1 cells with 'extramedullary' HSC/MPP tier1 cells. FDR <0.05. **B)** Split violin plots of selected histone genes, significantly upregulated (FDR<0.05 by EdgeR) in 'pan-tissue' (orange) HSC/MPP tier1 cells compared to 'extramedullary' (blue) HSC/MPP tier1 cells. Solid lines indicated the median in all violin plots. **C)** Phenotypic HSC/MPPs ($CD19^-$ $CD34^+$ $CD38^-$ $CD45RA^-$) from BM and spleen of the same paired donors as used for 10X RNA-seq were single-cell sorted and transcriptome was sequenced using the Smart-seq2 protocol. Bar graphs show population-specific signatures (left, (Laurenti et al., 2013, 2015)) and lineage-priming modules (right, (Velten et al., 2017)) from pre-ranked GSEA (FDR <0.05) comparing BM and spleen HSC/MPPs. **D-F:** single cell MEM differentiation assay performed with single HSC/MPP from BM or spleen. **D)** Colony efficiency calculated as the number of positive colonies (for definition see methods) as fraction of the number of seeded single cells. **E)** Bar graphs of the colony composition separated by sample. **F)** Fraction of Ery-containing colonies. Each dot represents one donor. SPL: spleen.

### SUPPLEMENTARY TABLES

**Table S1. Information on the deceased organ donors used in this study**. Tissues from the donors with the internal ID KSP11 and KSP23 were used for the 10x scRNA-seq and Smart-seq2 analyses and are referred to as ‘donor1’ and ‘donor2’, respectively (highlighted in green in the table).

**Table S2. Numbers of cells submitted for scRNA-seq, and after first steps of quality control prior deeper bioinformatics analyses.** The numbers of HSPCs loaded onto the 10x chip as well as the cell recovered of sequencing as determined with the Cell Ranger software are shown (tab1). A second tab lists the number of single HSC/MPPs submitted for RNA-seq with the Smart-seq2 protocol, as well as the number of cell transcriptomes passing quality control.

**Table S3. Absolute and relative proportions of cells within the 26 Louvain clusters** (tab1-2) in the Seurat-aligned dataset; the 6 main HSPC groups as defined in **Figure 1B** and shown in **1G, 2A, S1C-D, S2A** (tab3); lineage-committed progenitors as shown in **Figure 2C, S2B** (tab4); and within the Ery/Meg/Baso/MC branch (tab5, relating to **Figure 3B, S3**). Green rows highlight the cell numbers and proportions received after combining both donors. Values for each individual donor are shown below.

**Table S4. Top100 marker genes for each Louvain cluster.** The most differentially expressed genes in each Louvain cluster relative to the remainder clusters was determined as described in the methods. The top 100 genes for each cluster are summarized, ranked from the highest to lowest score of the Wilcoxon rank sum test.

**Table S5. Most frequently projected cluster per phenotypically isolated HSC/MPP.** Smart-seq2 single-cell transcriptomes of phenotypic HSC/MPPs were projected onto the HSPC landscape obtained by 10x scRNA-seq (see methods). Table summarized the most frequently projected cluster for each individual Smart-seq2 cell, considering its nearest 15 cell projections (tab1). Tab2 shows the same data, but grouped into the 6 main HSPC groups as defined in Figure 1B.

**Table S6. Transcriptome based cell cycle assignment.** Cell cycle phase scores were computed for each single cell as described in the methods. Table summarises the absolute number of cells from each cluster assigned to $G_1$, S and $G_2$-M phases and the relative proportion of cells from these clusters being in the respective cell cycle phases. Data was determined combining all three tissues (tab1) or individually for BM, PB and spleen (tab2).

**Table S7. Differential gene expression comparing BM MEP1 (cluster11 with splenic MEP1.** Results are shown as being up-/down-regulated in BM compared to spleen as reference tissue.

**Table S8. Results of genes set enrichment analysis in BM and splenic MEPs.** GSEA testing for enrichment of C2 curated gene sets was performed on pre-ranked differential gene expression lists from MEP1 (cluster11, tab1), MEP2 (cluster18, tab2) and MEP3 (cluster9, tab3) comparing BM (highest rank) and spleen (lowest rank). Only gene sets with FDR<0.05 are shown. Selected pathways from this list are shown in **Figure 3E**.

**Table S9. Differential gene expression comparing EryPs (cluster4) from BM with splenic MEP1.** Results are shown as being up-/down-regulated in BM compared to spleen, which was used as reference group.

**Table S10. List of NSG mice that were transplanted with spleen Lin- cells.** Human chimerism in the BM of recipient mice was analysed 14 weeks after transplantation.

**Table S11. Number of BM and spleen derived HSC/MPP tier1 cells within the 'extramedullary' and 'pan-tissue' clusters.**

**Table S12. Differential gene expression comparing cells belonging to the 'pan-tissue' and 'extramedullary' HSC/MPP Tier1 clusters.** Results are shown as being up-/down-regulated in extramedullary Tier1 HSC/MPPs compared to 'pan-tissue' Tier1 cells, which were used as reference group.

**Table S13. Gene set enrichment in 'pan-tissue' and 'extramedullary' HSC/MPP Tier1.** Gene set enrichment results testing for population-specific signatures (Laurenti et al., 2013, 2015) (tab1), lineage-priming signatures (Velten et al., 2017) (tab2) and MSigDB C2 curated gene sets (tab3) in extramedullary HSC/MPP Tier1 (highest rank) and 'pan-tissue' Tier1 cells (lowest rank). For the enrichment results of C2 gene sets, a cut-off of FDR<0.05 was used. Selected pathways from this list are shown in **Figure 4J**.

**Table S14. Gene set enrichment in BM and spleen phenotypic HSC/MPPs single-cell sequenced with the Smart-seq2 protocol.** Enrichment for population-specific signatures

(Laurenti et al., 2013, 2015) (tab1), lineage-priming signatures (Velten et al., 2017) (tab2) was tested.

**Table S15. Differential gene expression comparing BM and spleen phenotypic HSC/MPPs single-cell sequenced with the Smart-seq2 protocol.** BM was used as reference tissue, and results are shown as being up-/down-regulated in spleen.

**Table S16. List of antibodies used in flow cytometry.**

| Antigen | Conjugate | Clone | Company | Cat # | Dilution |
|---|---|---|---|---|---|
| CD3 | APC/Cy7 | HIT3a | BioLegend | 300318 | 1 : 100 |
| | FITC | HIT3a | BD Pharmingen | 555339 | 1 : 500 |
| CD5 | PE | UCHT2 | BD Pharmingen | 561897 | 1 : 300 |
| CD7 | BV421 | M-T701 | BD Horizon | 562635 | 1 : 100 |
| CD10 | APC | HI10a | BioLegend | 312210 | 1 : 100 |
| | BV421 | HI10a | BD Horizon | 562902 | 1 : 100 |
| CD11b | APC/Cy7 | ICRF44 | BioLegend | 301342 | 1 : 300 |
| CD14 | PE-Cy7 | M5E2 | BioLegend | 301814 | 1 : 1000 |
| CD15 | BV421 | MC-480 | BioLegend | 125614 | 1 : 200 |
| CD19 | Alexa Fluor 700 | HIB19 | BioLegend | 302226 | 1 : 300 |
| | BV785 | SJ25C1 | BioLegend | 363028 | 1 : 600 |
| | FITC | HIB19 | BioLegend | 302206 | 1 : 200 |
| CD33 | APC | P67.6 | BD Pharmingen | 345800 | 1 : 200 |
| CD34 | APC-Cy7 | 581 | BioLegend | 343514 | 1 : 100 |
| CD36 | APC | 5-271 | BioLegend | 336208 | 1 : 300 |
| CD38 | PE/Cy7 | HIT2 | BioLegend | 303516 | 1 : 100 |
| | BV510 | HIP8 | BD Horizon | 563250 | 1 : 200 |
| | FITC | HIP8 | BioLegend | 303704 | 1 : 1000 |
| CD45 | BV510 | HI30 | BioLegend | 304036 | 1 : 500 |
| | PE-Cy5 | HI30 | BioLegend | 304010 | 1 : 300 |
| | BV605 | 2D1 | BioLegend | 368524 | 1 : 100 |
| CD45RA | Alexa Fluor 700 | HI100 | BioLegend | 304120 | 1 : 300 |
| | FITC | HI100 | BD Pharmingen | 555488 | 1 : 100 |
| | PE | HI100 | BioLegend | 304108 | 1 : 200 |
| CD49f | PE-Cy5 | GoH3 | BD Pharmingen | 551129 | 1 : 100 |
| CD56 | APC | HCD56 | BioLegend | 318310 | 1 : 200 |
| CD90 | APC | 5E10 | BD Pharmingen | 559869 | 1 : 100 |
| | PE | 5E10 | BioLegend | 328110 | 1 : 50 |
| | FITC | 5E10 | BioLegend | 328108 | 1 : 50 |
| CD71 | FITC | CY1G4 | BioLegend | 334104 | 1 : 1000 |
| | PerCP/Cy5.5 | CY1G4 | BioLegend | 334113 | 1 : 800 |
| CD117 (c-Kit) | BV650 | 104D2 | BioLegend | 313222 | 1 : 100 |
| CD135 | PE | BV10A4H2 | BioLegend | 313305 | 1 : 50 |
| CD370 (CLEC9A) | PE | 8F9 | BioLegend | 353804 | 1 : 75 |
| GlyA | PE | GA-R2 (HIR2) | BD Pharmingen | 340947 | 1 : 1000 |
| | BV421 | GA-R2 (HIR2) | BD Horizon | 562938 | 1 : 1000 |
| KI-67 | FITC | B56 | BD Pharmingen | 556026 | 1 : 50 |
